# Circulating microRNAs reveal egg-brain crosstalk and a brain-specific microRNA linked to the onset of the next reproductive cycle in iteroparous salmonids

**DOI:** 10.64898/2025.12.22.695901

**Authors:** Mariana Roza de Abreu, Emilie Cardona, Camille Lagarde, Leo Milhade, Violette Thermes, Julien Bobe

**Affiliations:** INRAE, LPGP, 35000 Rennes, France; INRAE, Univ. Pau & Pays Adour, E2S UPPA, NUMEA, 64310, Saint-Pée-sur-Nivelle, France; IRISA, INRIA, CNRS, Université de Rennes 1, 35000, Rennes, France

**Keywords:** biomarker, egg quality, miR-139, miR-202, miR-457a, miR-135c, iteroparity

## Abstract

Mechanisms regulating the transition between two consecutive reproductive cycles are complex and remain poorly understood, mostly because they involve a dialog between the ovary and the central nervous system that is difficult to disentangle. In rainbow trout (*Oncorhynchus mykiss*), an iteroparous species spawning every year, removal of the eggs from the body cavity was used as a switch to trigger the onset of the next reproductive cycle. Changes in circulating miRNAs (c-miRNAs) levels in blood plasma and ovarian fluid were then monitored over time. Upon removal of the eggs from the body cavity we observed the dramatic down regulation of the blood plasma levels of a single c-miRNA (miR-139-5p) that is predominantly expressed in the brain. In contrast, very distinct c-miRNAs profiles were observed in blood plasma when eggs are retained in the body cavity. Among plasma c-miRNAs showing dynamic changes with egg retention, miR-135c is strongly expressed in the brain and pituitary, while miR-457a is predominant in the postovulatory ovary. In addition, egg retention in the body cavity triggers a dramatic drop in ovarian fluid levels of miR-202-5p, a miRNA known to regulate egg production in fish. Our observations reveal that the transition between two successive reproductive cycles involves a crosstalk between the eggs and the central system and that a single miRNA, miR-139, predominantly expressed in the brain, is associated with the onset of the next reproductive cycle. We identified possible miR-139-5p functional targets in rainbow trout and other iteroparous species that have been lost in semelparous salmonids. Our results offer new research perspectives to better understand the mechanisms triggering the next reproductive cycle in iteroparous fish species, including post-transcriptional regulations by miR-139 in the brain.

## Introduction

In contrast to mammals, ray-finned fish species exhibit continuous female germ cell proliferation throughout their adult life [1]. Thus, most fish species have the ability to spawn multiple times. Even though different types of ovarian dynamics have been reported, from synchronous to asynchronous oocyte growth and subsequent ovulation [1], the release of the eggs from the ovarian or body cavity into the water at the time of spawning represents a checkpoint regulating the switch from one reproductive cycle to the next. The post-ovulatory period is also critical for reproductive success as ovulated oocytes (*i.e.* eggs) exhibit a progressive, and usually rapid, decrease of their ability to be fertilized [2]. During this period, eggs are held either the ovarian lumen or the body cavity, depending on the species, in a biological fluid referred to as ovarian fluid [3]. However, despite the importance of the post-ovulatory period (*i.e*. between ovulation and spawning) in transitioning from one reproductive cycle to the next, the mechanisms at stake remain elusive.

Several reasons explain why our knowledge of this key phase between two reproductive cycles remains limited, including the difficulty to integrate the crosstalk between peripheral and central regulations, and associated molecular mechanisms. To go beyond the state of the art and explore the transition between two reproductive cycles with a fresh look, we proposed to use circulating miRNAs. These short (∼ 22-nt) non-coding RNAs involved in the post-transcriptional regulation of gene expression [4] have recently emerged as highly promising non-invasive biomarker molecules that can reflect physio pathological status due to their high stability and presence in most, if not all, biological fluids. In humans, c-miRNAs have been proposed as non-invasive biomarkers of many pathologies and their relevance to serve as diagnostic or prognostic tools has been established [5–7]. The levels of specific miRNAs in biological fluids, including blood plasma, is therefore highly informative to reveal the activation of specific mechanisms in various organs as recently shown in fish [8].

To investigate the transition from one reproductive cycle to the next, we used rainbow trout (*Oncorhynchus mykiss*), a species with known c-miRNA repertoire [8]. In this iteroparous species, the post-ovulatory period is not only characterized by a slow and progressive decrease in gamete quality [2], but is also important for the recruitment of the next batch of ovarian follicles into the reproductive cycle. Several lines of evidence have shown that the release of the eggs from the body cavity triggers a switch in hormonal levels from LH, the hormone regulating final oocyte maturation and ovulation, to FSH that regulates oogonial proliferation and follicular recruitment into the reproductive cycle [9]. These observations indicate that the release of the eggs from the body cavity represents a checkpoint in the reproductive cycle that regulates the onset of the next wave of follicular recruitment into the reproductive cycle. However, besides the switch in hormonal levels, very little is known about how this transition from one cycle to another is regulated, especially at the molecular level.

To go beyond endocrine regulations, we monitored dynamic changes in c-miRNA levels in blood plasma and ovarian fluid, the biological fluid in which unfertilized eggs are held upon ovulation, over time, and in response to the removal of the eggs from the body cavity that was used a switch to trigger the onset of the next reproductive cycle. Upon removal of the eggs from the body cavity, we observed the dramatic down regulation in blood plasma levels of a single c-miRNA (miR-139-5p) that is highly and predominantly expressed in the brain. This event is likely to represent a molecular switch corresponding to the onset of the next reproductive cycle. Our conclusions are further supported by the very distinct and dynamic c-miRNAs profiles observed when eggs are retained in the body cavity. Among c-miRNAs exhibiting marked dynamic changes in blood plasma associated with egg retention were mR-135c and miR-457a. While miR-139 and miR-135c exhibit a strong predominant expression in the brain (and in the pituitary in the case of miR-135c), miR-457a is predominantly expressed in the post-ovulatory ovary. These expression patterns appear to be conserved in other teleost fish species including zebrafish (*Danio rerio*) and medaka (*Oryzias latipes*). In addition, egg retention in the body cavity triggered a dramatic drop in ovarian fluid levels of miR-202-5p, a miRNA known to regulate egg production in medaka [10, 11]. While these observations argue in favor of a functional role played by miR-202 in regulating post-ovulatory ovarian functions, we also showed that miR-202-5p levels in ovarian fluid can be used as an efficient non-invasive marker of egg quality.

Together, our observations clearly demonstrate that holding or removing the eggs from the body cavity – an event leading to the onset of the next reproductive cycle – is associated with very distinct physiological processes that are clearly reflected by blood plasma c-miRNA profiles. We show that the transition between two successive reproductive cycles involves a crosstalk between the ovary and the central system and that a single miRNA, miR-139, predominantly expressed in the brain is associated with the onset of the next reproductive cycle. We identified possible miR-139-5p functional targets in rainbow trout and other iteroparous salmonid species that have been lost in semelparous salmonids. Our results offer new research perspectives to better understand the mechanisms triggering the next reproductive cycle in iteroparous fish species, including post-transcriptional regulations by miR-139 in the brain.

## Results

### Egg removal triggers the onset of the next reproductive cycle

In order to demonstrate that egg removal from the body cavity triggered the onset of the next reproductive cycle, we monitored ovarian follicle size and number on histology sections. While the average follicular diameter was not significantly different, we observed a significant increase in the number of follicles, after three weeks, when eggs when removed from the body cavity at ovulation (Fig 1A). While we did not observe any difference in mean follicular diameter (Fig 1B), the difference in follicle count was especially marked for previtellogenic follicles exhibiting an observed size range of 300-600 μm on histological sections (Fig 1C).

**Figure 1:**
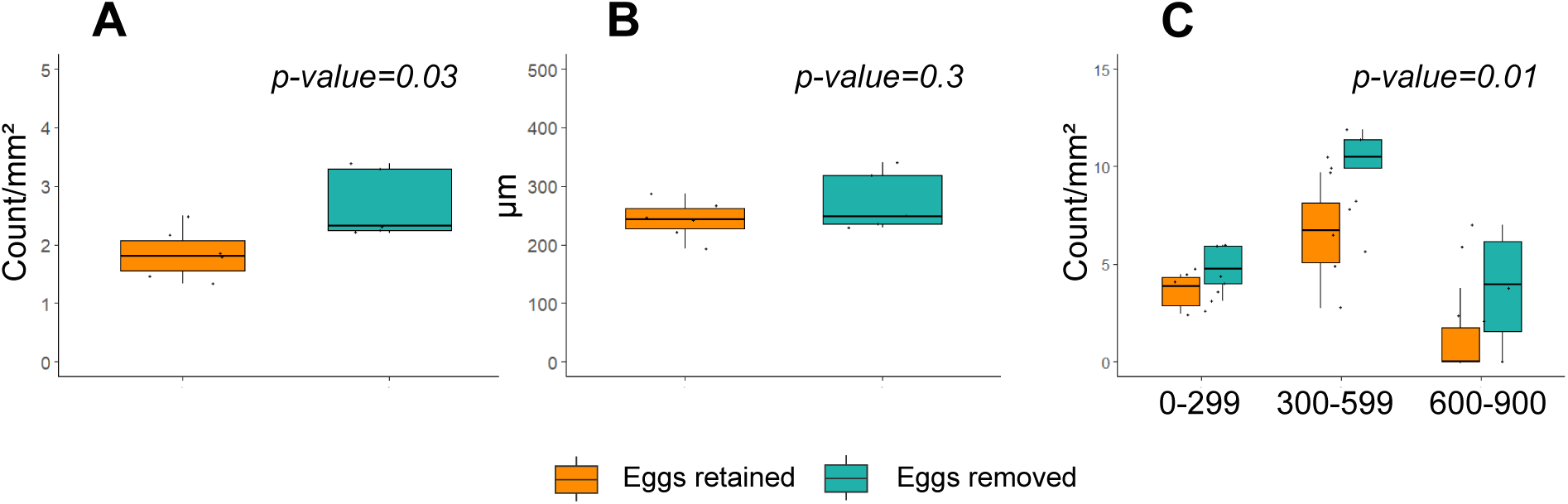
Impact of egg removal or retention of follicular dynamics in the ovary. **A**: Total number of ovarian follicles on ovarian sections. **B**: Mean ovarian follicular diameter. **C**: Number of ovarian follicles of different size classes.

### Blood plasma and ovarian fluid exhibit distinct, yet overlapping, c-miRNA repertoires

Among the 354 annotated rainbow trout miRNAs [8], 221 were detected above a threshold of 10 reads per million reads, on average, either in blood plasma or ovarian fluid. Among these 221 miRNAs, 176 (80%) were present in both blood plasma and ovarian fluid, while 22 (10%) were detected only in plasma and 23 (10%) were detected only in ovarian fluid (Fig 2A). Among the 10 most abundant miRNAs in ovarian fluid and blood plasma, six were common to both fluids (miR-451-5p, let-7a-5p, miR-21-5p, miR-92a-3p, miR-22a-1-3p and let-7e-5p). In blood plasma miR-150-5p, miR-16b-5p, miR-26a-5p, and miR-100-5p were the four most abundant miRNAs. In ovarian fluid miR-202-5p, miR-146a-5p, miR-125b/c-5p, and miR-30d-5p were the four other most abundant miRNAs (Fig 2B). Despite some similarities between both fluids, the ovarian fluid and blood plasma miRNAomes could be clearly discriminated. The Principal Component Analysis (PCA) showed that the overall c-miRNA profiles differ between blood plasma and ovarian fluid at ovulation (Fig 3A) and 21 days after ovulation (Fig 3B). At both stages, the differences between the two biological fluids investigated appear to be driven, at least in part, by a limited number of specific c-miRNAs that are likely to be of physiological relevance. However, the identity of the miRNAs discriminating the two fluids was different at ovulation and 21 days after ovulation (Fig 3).

**Figure 2:**
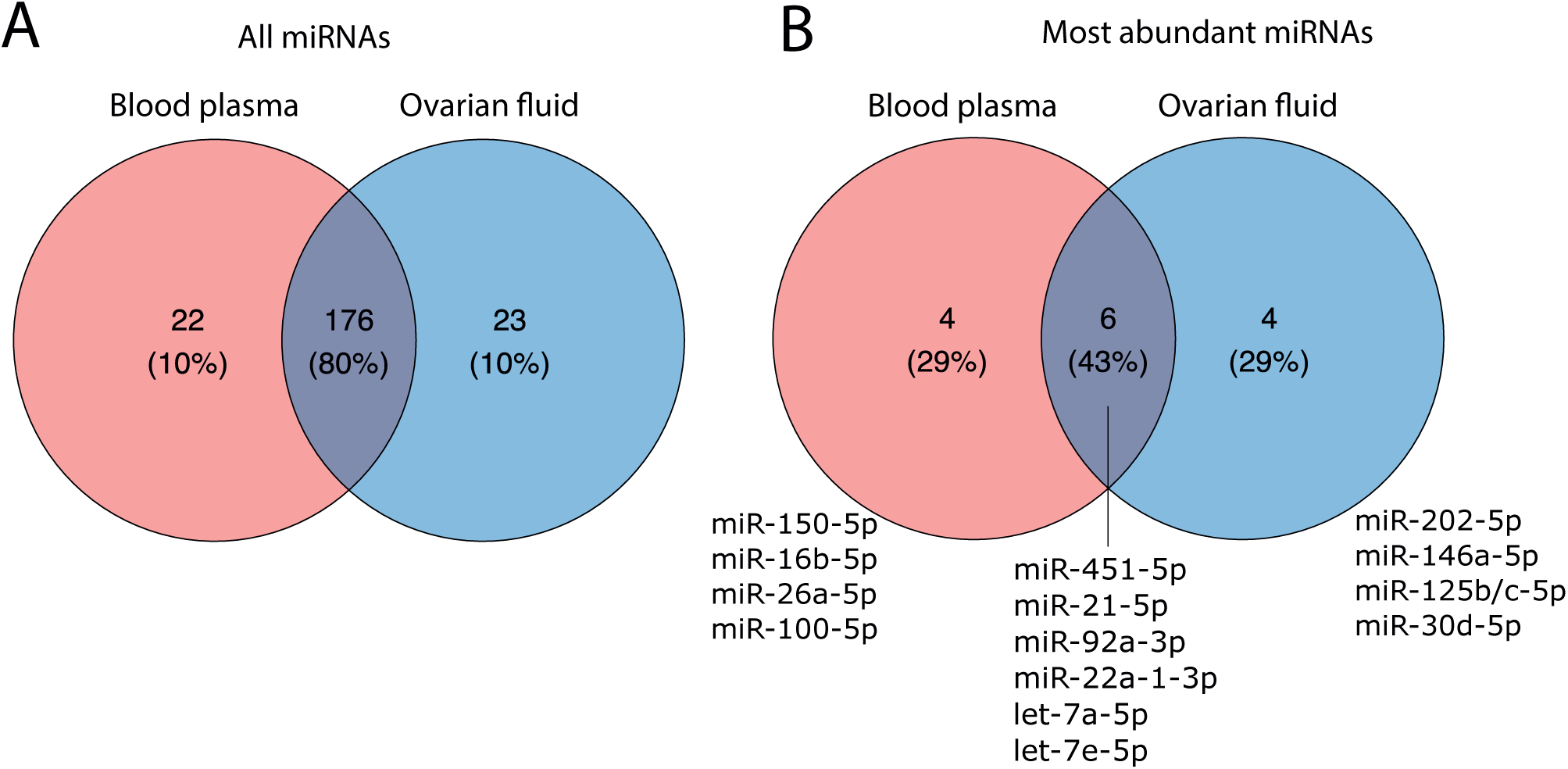
Circulating miRNA repertoire in blood plasma and ovarian fluid. A specific miRNA was considered expressed in a sample when its averaged normalized abundance exceeded 10 RPM (reads per million reads). (**A**) Venn diagram of all miRNAs detected in blood plasma and ovarian fluid. (**B**) Top 10 most expressed miRNAs in blood plasma and ovarian fluid.

**Figure 3:**
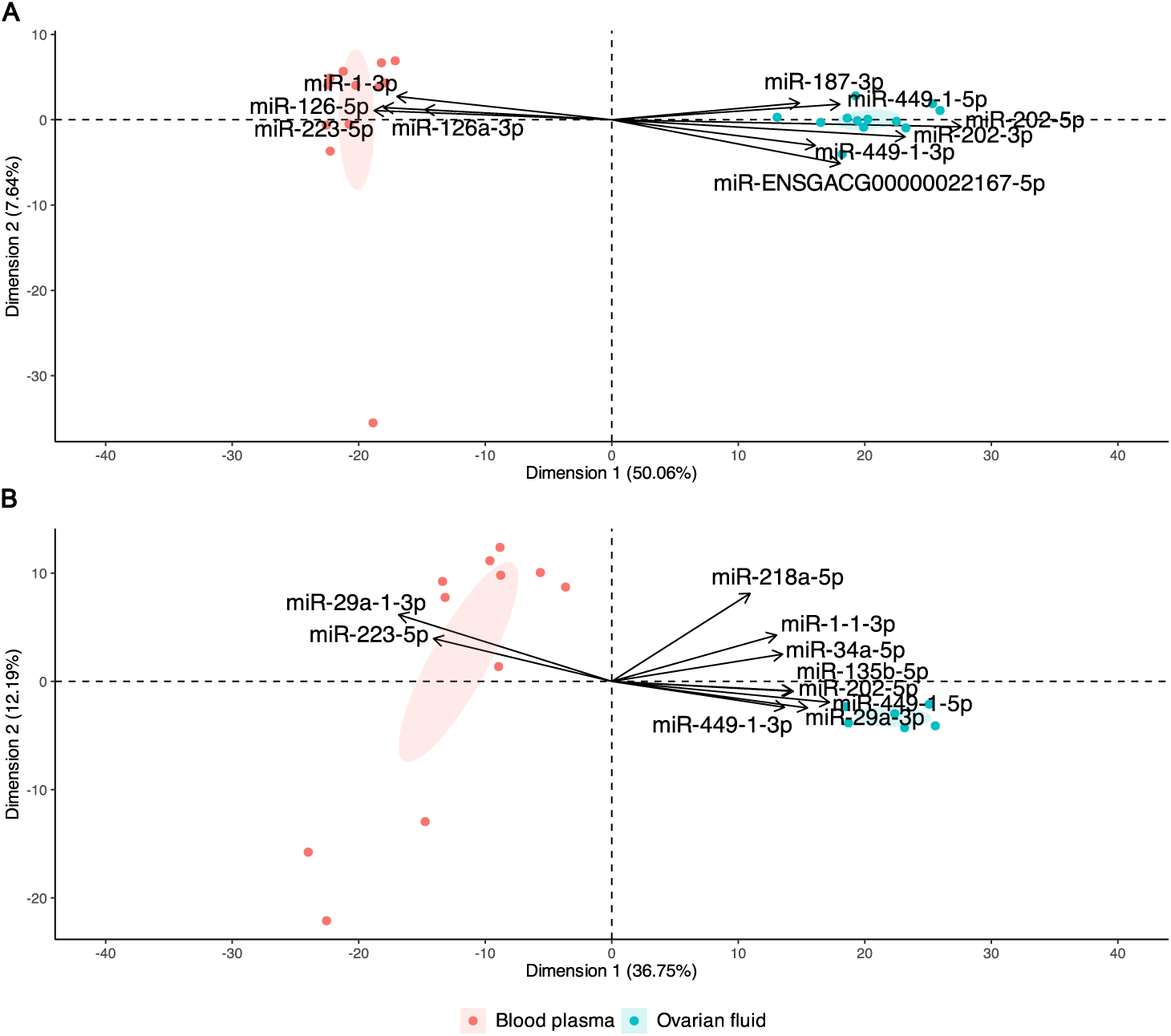
A: PCA analysis of normalized RPM counts using all samples at ovulation. **B**: PCA analysis of normalized RPM counts using all samples at 21 days post-ovulation. PCAs were centered but not scaled and computed from normalized miRNAs counts (RPM) supplied by Prost! [12]. Ellipses represent 95% confidence intervals and were drawn for conditions represented by at least three individuals.

### A single c-miRNA is associated with the onset of the next reproductive cycle

During the post-ovulatory period, c-miRNA profiling revealed major dynamic changes over time in blood plasma when eggs were retained in the body cavity (Fig 4A, Supplementary data file 1). In sharp contrast with these observations, removing the eggs from the body cavity upon ovulation, yielded very different results, with a single c-miRNA exhibiting a significant down-regulation during the post-ovulatory period (Fig 4B, Supplementary data file 2). Together, our observations clearly demonstrate that holding or removing the eggs from the body cavity – an event leading to the onset of the next reproductive cycle – is associated with very distinct physiological processes that are clearly reflected by blood plasma c-miRNA profiles.

**Figure 4:**
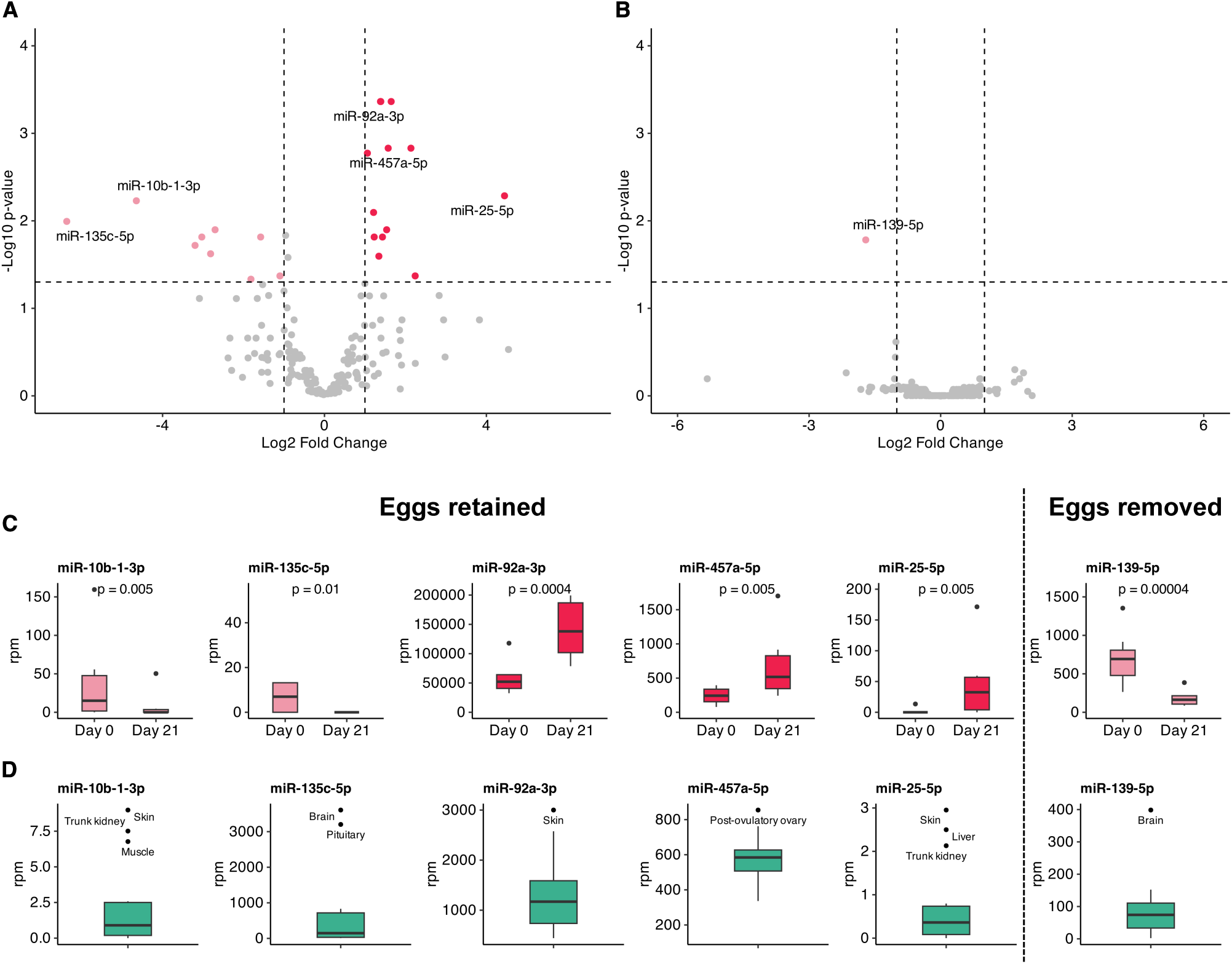
Blood plasma c-miRNA levels when eggs are retained or removed from the body cavity between ovulation (Day 0) and 21 days post-ovulation (Day 21). Differentially abundant c-miRNA when eggs are retained in the body cavity for 21 days (**A**) or removed from the body cavity at ovulation (**B**). Volcano plots show differentially abundant c-miRNAs (Log2 fold change > 2, Benjamini-Hochberg adjusted p value < 0.05). Up-regulated and down-regulated c-miRNAs at Day 21 relative to Day 0 are indicated in red and pink, respectively. **C:** Boxplots showing the expression levels of specific c-miRNAs in blood plasma at ovulation (D0) and 21 days after ovulation (D21). **D**: Boxplots showing miRNA expression in different organs. The organs exhibiting the maximum expression are indicated.

When eggs are removed from the body cavity upon ovulation, we observed a dramatic and highly significant down-regulation of a single c-miRNA (miR-139-5p) in blood plasma (Fig 4C), while all other detected c-miRNAs remain unaffected. This pattern is not observed when eggs are kept in the body cavity throughout the period. To investigate the possible organ of origin of miR-139-5p, we categorized c-miRNAs based on the organ in which they exhibited the highest expression. This analysis investigated a set of 13 organs (brain, gills, head kidney, heart, intestine, liver, muscle, post-ovulatory ovary, pituitary, skin, spleen, stomach, trunk kidney) and unfertilized eggs (Supplementary data file 3) using dataset that we had previously generated [8, 13]. Our data showed that miR-139-5p was highly and predominantly expressed in the brain (Fig 4D). Together these observations, including the unicity and amplitude of the response observed, strongly suggest that miR-139-5p is associated with the switch between two successive reproductive cycles triggered by the release of the eggs from the body cavity. Our data suggest that this switch involves, at least in part, post-transcriptional regulations by miR-139-5p in the brain. Studies in mammals have shown that miR-139-5p plays an important role in brain functions and exhibits a differential expression in response to stress or in specific pathologies, including depression, Parkinson’s disease, epilepsy, brain tumor [14, 15]. This miRNA has also been identified as a relevant blood microRNA biomarker of resilience or vulnerability to stress [16]. In consistency with these results, expression data from the FishmiRNA database [17] show a strong and highly predominant expression of miR-139a-5p in the brain in zebrafish (*Danio rerio*) and medaka (*Oryzias latipes*).

In contrast, when eggs were retained in the body cavity, we identified 13 up-regulated c-miRNAs in blood plasma over the post-ovulatory period, all exhibiting a marked up-regulation (Log2 FC > 1). We also identified 11 down-regulated c-miRNAs, including 9 c-miRNAs exhibiting a marked (Log2 FC >1) down-regulation (Fig 4A). Among the top 5 c-miRNAs exhibiting the most significant differences and highest fold-change were two down-regulated (miR-10b-1-3p and miR-135c-5p) and three up-regulated (miR-92a-3p, miR-457a-5p, and miR-25-5p) c-miRNAs (Fig 4C). We observed that miR-457a-5p was predominantly expressed in the post-ovulatory ovary (Fig 4D). These observations suggest an over expression of this miRNA in the ovary over time during the post-ovulatory period. When investigating expression patterns of miR-457a-5p in the FishmiRNA database [17], we observed a strong and predominant ovarian expression of this miRNA in other teleost species, including zebrafish (*Danio rerio*) and medaka (*Oryzias latipes*). Together, these observations suggest an important role of miR-457a-5p in regulating ovarian functions when eggs are held in the body cavity. This hypothesis is further supported by the lack of differences in circulating miR-457a-5p levels over time during the post-ovulatory period when eggs are removed from the body cavity (Fig 4B). To our knowledge, the importance of miR-457a in ovarian functions was previously unsuspected.

In contrast to miR-457a-5p, other miRNAs exhibiting marked fold-change and highly significant differences are predominantly expressed in non-reproductive organs. Among them, miR-135c-5p was the c-miRNA exhibiting the most pronounced down-regulation during the post-ovulatory period (Fig 4A). This miRNA was predominantly expressed in brain and pituitary (Fig 4D). In zebrafish and medaka, miR-135c-5p is also highly predominantly expressed in the brain, even though it appears expressed, at much lower levels, in reproductive organs. Together, the dynamics of expression and tissue distribution of miR-135c-5p suggest an important role, at the central level, in regulating physiological processes at stake while eggs are held in the body cavity. The marked down-regulation during the post-ovulatory period suggests suppression that the post-transcriptional repression of miR-135c-5p targets is required. We hypothesize that these target genes could be involved in either regulating ovarian functions and egg preservation, or in blocking the onset of the next reproductive cycle.

In addition to miR-457a and miR-135c-5p, three other miRNAs (miR-10b-1-3p, miR-92a-3p and miR-25p-5p) exhibited a marked differential regulation during the post-ovulatory period (i.e., between ovulation and egg removal). While miR-10b-1-3p was down-regulated, miR-25-5p and miR-92a-3p were up-regulated, the later exhibiting a dramatic up-regulation. As indicated above, miR-92a-3p is among the 10 most abundant miRNAs in blood plasma. In rainbow trout, miR-92a exhibits a predominant expression in skin in the subset of samples (brain, gills, head kidney, heart, intestine, liver, muscle, post-ovulatory ovary, pituitary, skin, spleen, stomach, trunk kidney, and unfertilized eggs) that we primarily investigated. However, miR-92a-3p is also expressed at significant levels in various organs, including the ovary. In this context, while it seems difficult to speculate on the specific role(s) of miR-92a during the post-ovulatory period, the expression levels in different organs and blood plasma, as well as the dramatic increased in blood plasma levels during the period, suggest a significant biological role, possibly at the ovarian level, that remains to be investigated.

In summary, the presence or absence of the eggs in the body cavity triggers major physiological changes with distinct outcomes in terms of blood plasma c-miRNA profiles. Removal of the eggs from the body cavity results in a dramatic decrease in blood plasma levels of a single c-miRNA (miR-139-5p) that is predominantly expressed in the brain and known to play important role in brain fucntions. In contrast, egg retention in the body cavity triggers a different physiological response involving miR-135c-5p, another brain predominant miRNA. Together, these observations indicate that the release of the eggs from the body cavity triggers specific mechanisms at the central level, including miRNA-mediated post-transcriptional regulation. We also demonstrate that egg retention in the body cavity is associated with dynamic changes of various c-miRNAs in blood plasma, including the up-regulation of miR-457a, a miRNA predominantly expressed in the post-ovulatory ovary whose role in ovarian functions was previously unsuspected.

### Egg retention in the body cavity is associated with major changes in ovarian fluid c-miRNAome

As indicated above, egg removal from the body cavity triggers a rapid transition between two consecutive cycles as suggested by previous observations on oogonial mitosis and dynamic changes in FSH and LH plasma levels, and confirmed by our observations. The importance of this phenomenon is highlighted by the marked differences in c-miRNA levels observed upon retention or removal of the eggs from the body cavity. However, the post-ovulatory period also plays a key role in the reproductive process. Salmonid fishes, unlike most other teleost fishes, have the ability to preserve egg viability for several days, even though long retention times are associated with a significant decrease of egg quality [2, 18, 19]. To delve deeper into ovarian regulations during this critical period we investigated c-miRNA levels in ovarian fluid that plays a key role in preserving egg viability. When comparing c-miRNA profiles in ovarian fluid between ovulation and 21 days post-ovulation, which is usually associated with severe egg quality defects due to post-ovulatory ageing [20], we identified 78 differentially expressed miRNAs including 27 down-regulated and 51 up-regulated miRNAs (Fig 5A, Supplementary data file 4). When focusing on c-miRNA exhibiting marked differences (Log2 FC >1 and adjusted p value < 0.05), we observed that miR-202-5p was highly expressed at ovulation and dramatically down-regulated at 21 days post-ovulation. The opposite pattern was observed for miR-15a-5p, miR-1-3p, miR-142-a-3p, and miR-144-5p, which were upregulated at 21 days post-ovulation compared to ovulation (Fig 5B). When investigating possible organ of origin of these c-miRNAs, we observed that miR-202-5p was overabundant in unfertilized eggs and post-ovulatory ovary. In contrast miR-1-3p was overabundant in muscle, heart, and stomach and miR-142-a-3p was found to be overabundant in trunk kidney (Fig 5C).

**Figure 5.**
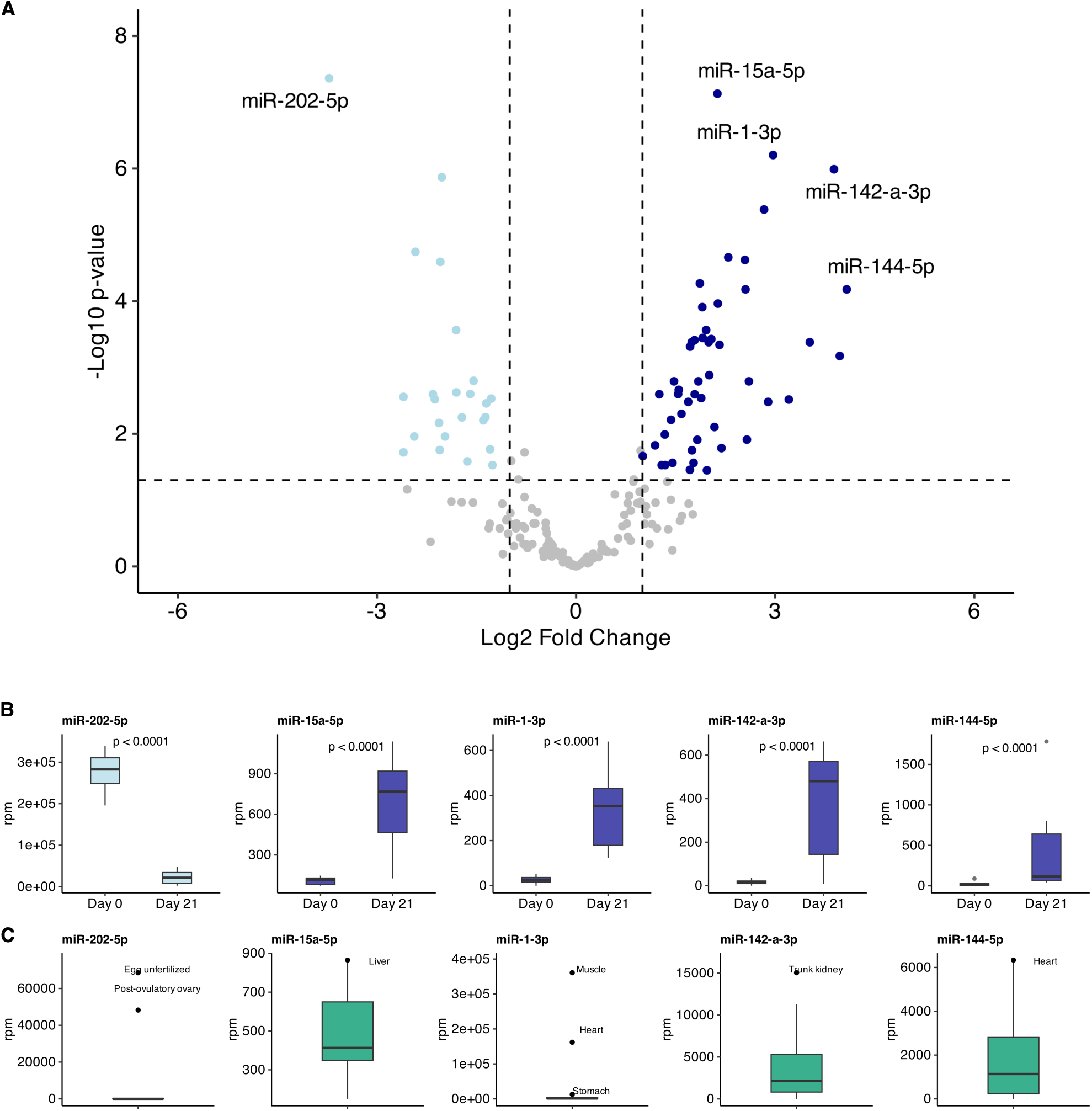
Ovarian fluid c-miRNA levels when eggs are retained in the body cavity. **A**: The volcano plot shows differentially abundant c-miRNAs (Log2 fold change > 2, Benjamini-Hochberg adjusted p value < 0.05) between ovulation (Day 0) and 21 days post-ovulation (Day 21). Up-regulated and down-regulated c-miRNAs at Day 21 relative to Day 0 are indicated in dark blue and light blue, respectively. **B:** Boxplots showing the expression levels of specific c-miRNAs in ovarian fluid at ovulation (D0) and 21 days after ovulation (D21). **C**: Boxplots showing miRNA expression in different organs. The organs exhibiting the maximum expression are indicated.

### Ovarian fluid c-miRNAs as non-invasive markers of egg quality

Small amounts of ovarian fluid can be non-invasively collected through gentle hand stripping. In this context, ovarian fluid c-miRNA could therefore be used as non-invasive markers. However, when investigating individual variability (Fig 6), we observed that only miR-1-3p and miR-202-5p exhibited non overlapping expression levels between ovulation (*i.e*., good quality eggs) and post-ovulation (*i.e*., low quality eggs). However, the difference in normalized counts of miR-1-3p between the ovulation samples exhibiting the highest values and the post-ovulation samples exhibiting the lowest values were rather limited (less than 2-fold). In contrast, very clear and marked differences were observed for miR-202-5p. For these reasons and also because levels in ovarian fluid are extremely high, we propose that miR-202-5p levels in ovarian fluid can be used as a biomarker to assess the drop of egg quality triggered by post-ovulatory ageing after ovulation.

**Figure 6.**
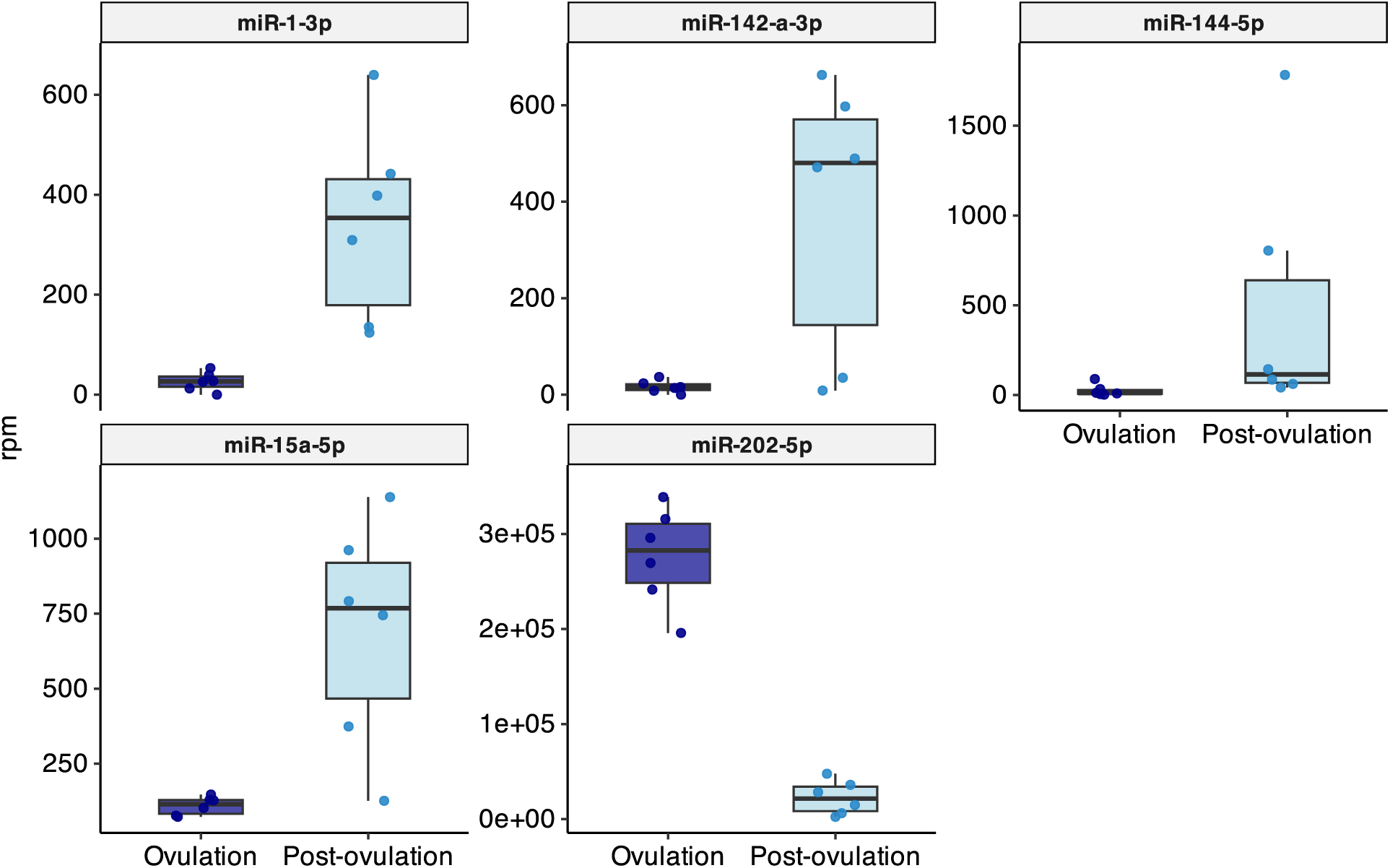
Ovarian fluid c-miRNA levels at ovulation (Day 0) and 21 days post-ovulation (Day 21), when eggs are retained in the body cavity. Individual values corresponding to different females are displayed.

### Possible miR-139-5p targets linked to iteroparity

Identifying miRNA targets is very challenging because the sequence recognized by the miRNA corresponds to 6-8 nucleotides matching the seed sequence [4] that can be found frequently in the 3’UTR sequence of many messenger RNAs. Thus, *in silico* target predictions can results in several hundreds or thousands of possible targets. However, several studies in animals, including fish [11], have shown that miRNAs can act *in vivo* through a single biological target to trigger a specific phenotype ([21, 22]). To refine target prediction, it is thus possible to include several features such as low variability in gene expression, the type of seed (6, 7 or 8 nucleotides), the spatiotemporal co-expression of the miRNA and the seed in a specific cellular context, the affinity of the miRNA for the target site, and the conservation of the seed [11]. In our study, miR-139-5p targets were predicted using the rainbow trout 3’UTRs available in Ensembl. This *in silico* analysis yielded 3,321 possible miR-139-5p targets, corresponding to 2,801 annotated rainbow trout genes (supplementary datafile 5). To identify relevant biological targets, we hypothesized that a target site involved in the onset of the next reproductive cycle was more likely to be conserved in iteroparous salmonid species, but lost (or at least less conserved) in semelparous salmonid species that reproduce only once. Similarly to Atlantic salmon (*Salmo salar*) and Brown trout (*Salmo trutta*), Rainbow trout (*Oncorhynchus mykiss*) is an iteroparous species that can spawn multiple times. In contrast, Pacific salmons such as coho salmon (*Oncorhynchus kisutch*) and Chinook salmon (*Oncorhynchus tshawytscha*) are semelparous salmonids that can only spawn once, then die. The loss of a specific target in coho salmon and Chinook salmon, which are phylogenetically closer to rainbow trout, and its conservation in brown trout and Atlantic salmon, which are more phylogenetically more distant, would support a link with iteroparity. Under this hypothesis, we identified 56 genes exhibiting a miR-139-5p target site in their 3’ UTR in iteroparous, but not semelparous, species (Fig. 7A, supplementary datafile 5). For 16 of those genes, the orthologous gene was missing from the dataset in at least one of the two semelparous species, while the 3’UTR was missing in at least one of the two semelparous species in 22 cases. We decided to focus on the remaining 23 genes for which a miR-139-5p target site was unambiguously present in iteroparous species while absent in semelparous species (Fig. 7A). To support the hypothesis of miR-139-5p target sites conserved in iteroparous species, we incorporated several features including the conservation of the target site in terms of location within the UTR and type of target site (6mer, 7mer-1a, 7mer-m8, 8mer), as well as the binding affinity for the target (Fig. 7A). We also incorporated existing RNA-seq data in rainbow trout from the AQUA-FAANG project [23] to monitor expression levels in different organs, under the assumption that the biological targets of miR-139, a brain-predominant miRNA, would also be expressed in the brain. Among the predicted targets exhibiting conserved high-affinity target sites, we noticed that several genes, including *dgkg*, *grid2*, *kcnip1b*, *slc24a1*, *vax1*, and *zfyve1* exhibited a significant expression in the brain (Fig. 7B). In the case of *dgkg*, *grid2*, *kcnip1b*, and *slc24a1*, the expression was even brain-predominant, which would be consistent with a major role in brain functions. Those genes are credible biologically relevant miR-139 targets that could be involved in regulating the onset of the next reproductive cycle.

**Figure 7.**
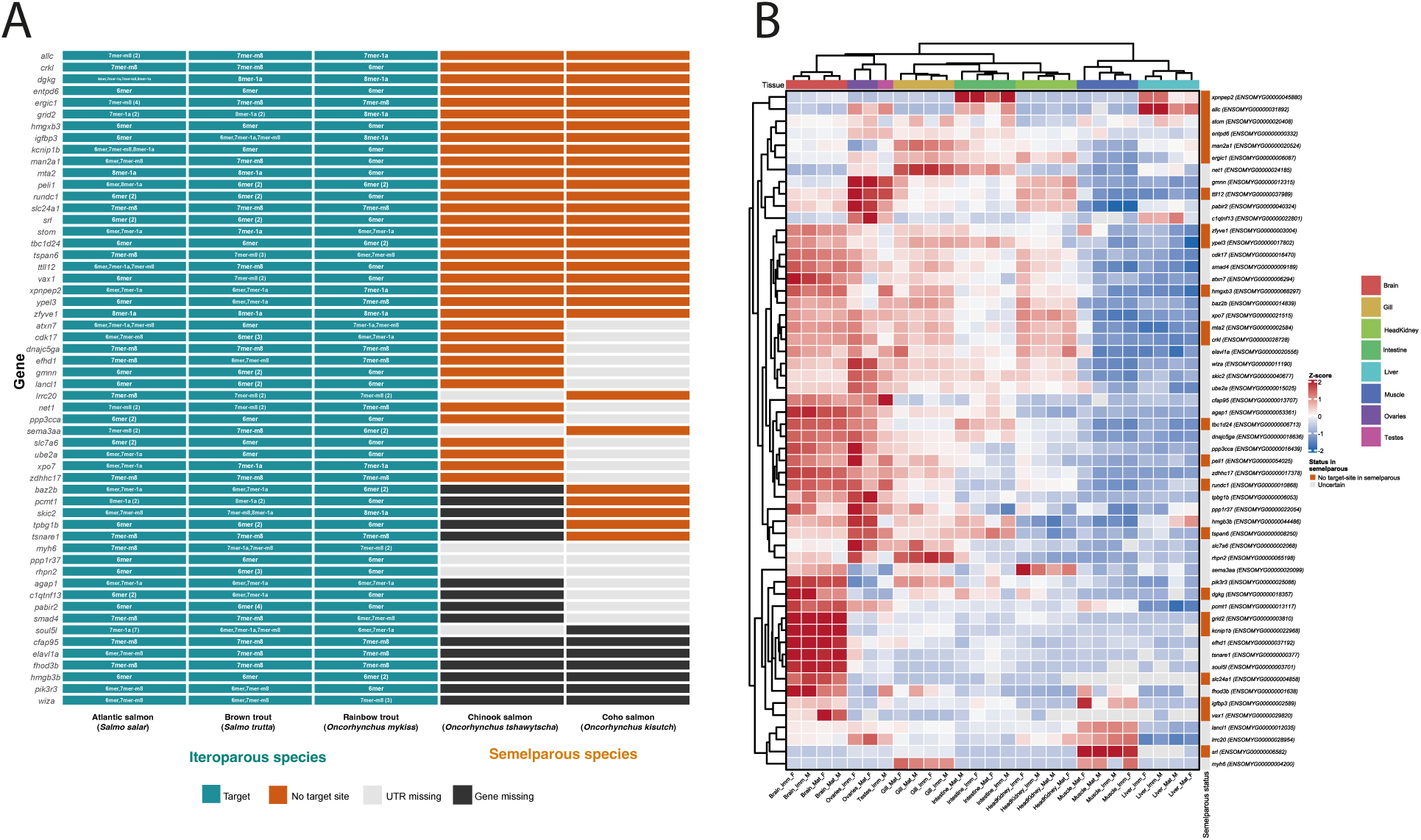
Possible miR-139-5p targets present in iteriparous, bur not semelparous, salmonid species (A). Expression levels of genes exhibiting a miR-139-5p binding site in their 3’UTR across various rainbow trout tissues and organs.

## Discussion

### A single brain-predominant miRNA linked to the switch between two reproductive cycles

In iteroparous salmonid species, the post-spawn period is usually associated with the onset of the next reproductive cycle. However, our understanding of the physiological and molecular mechanisms regulating the switch from one reproductive cycle to the next remains limited. In the wild, post-spawned individuals are energetically depleted because investment in both reproductive tissue and spawning behavior is energetically demanding [24]. For this reason, the role of the metabolic status during the post-spawn period has been investigated with a focus on endocrine factors [25, 26]. In rainbow trout, a pioneer work has showed that removing the egg from the body cavity triggered a rise in follicle stimulating hormone (FSH) plasma levels, one week after egg removal from the body cavity [9]. The post-spawn period is also associated with a rise in E2 synthesis capacity in the ovary, consistent with follicular development in the ovary [27]. However, the mechanisms triggered by the removal of the egg from the body cavity remain elusive both in the ovary and the central nervous system (CNS). In the present study, we have used a novel approach and monitored c-miRNAs in both blood plasma and ovarian fluids (in which eggs are held in the body cavity prior to spawning). Monitoring c-miRNAs in biological fluid is powerful to detect molecular signatures of specific biological processes or physiological states, as previously demonstrated in rainbow trout [8] and many animal species.

Here we show that egg removal from the body cavity drives changes in the blood plasma levels of a single c-miRNA, miR-139-5p. This observation is in sharp contrast with the dynamic changes in many blood plasma c-miRNAs observed when eggs are retained in the body cavity. In rainbow trout, as in other teleost species such as zebrafish and medaka [17], miR-139 is predominantly expressed in the brain. Together, these observations strongly suggest that the onset of the next reproductive cycle, triggered by removal of the eggs from the body cavity, is associated with the post-transcriptional regulation of specific target(s) by a single miRNA in the brain. The differences in c-miR-139-5p levels in blood plasma are observed as early as 21 days following egg removal, suggesting that miR-139-associated regulations are occurring relatively soon (i.e. less than three weeks) after egg removal.

This timing is consistent with the increase in FSH plasma levels that can be observed 7 days after egg removal [9] and the differences in follicle size that are already observed 3 weeks after ovulation in the present study. The hypothesis of a pivotal role played by miR-139-5p in the onset of the next reproductive cycle is further supported by the very distinct profiles in c-miRNA blood plasma profiles observed when eggs are retained in the body cavity, including miRNAs predominantly expressed in the CNS (miR-135c-5p) and the ovary (miR-457a-5p). More specifically, the observed drop in miR-135c-5p and the rise in miR-457a-5p suggest that blocking the onset of the next reproductive cycle also involves miRNA-mediated gene regulations in the CNS and the ovary, respectively. Given the profile of miR-135c-5p during the post-ovulatory ovary, it is also possible that this miRNA plays a role in the CNS during spawning, rather than in the physiological processes associated with egg retention in the body cavity. Together, our results reveal a crosstalk between the ovary and the CNS involving miRNA-mediated gene regulations that were previously unsuspected. They also suggest the importance of post-translational regulation of gene expression by a limited number of miRNAs in the onset of the next reproductive cycle in an iteroparous species.

Our results highlight the power of monitoring blood plasma c-miRNAs by revealing marked differences of a very limited number of biologically relevant miRNAs, tightly associated with spawning, and predominantly expressed in key organs including post-ovulatory ovary (miR-457a and miR-202), brain (miR-139 and miR-135c) and pituitary (miR-135c). While the importance of miR-202 for reproduction is documented in fish, the role of miR-457a, miR-139, and miR-135 was previously unsuspected. Identifying these new players using different approaches such as miRNA profiling in different organs would have been labor-intensive and would have precluded repeated sampling from the same individuals. However, despite the originality of our results, further investigations are needed to confirm the differential expression of identified miRNAs in the brain or in the ovary upon egg retention or removal. In the long term, identification of miR-202-5p, miR-139-5p, miR-135c-5p, and miR-457a-5p biological targets in their organs of expression would provide further insights into the molecular processes involved. However, going beyond the in silico prediction of putative miRNA targets to identify real biological targets remains a major challenge [11]. The identification of miRNA phenotypical targets (*i.e*. that contribute to the phenotypic outcome) is methodologically challenging, especially in non-model species with long generation time in which genome editing is labor intensive.

Among credible miR-139 targets, *dgkg*, *grid2*, *kcnip1b*, and *slc24a1* are of particular interest due to their predominant expression in the brain, including in sexually mature females. Because the post-transcriptional regulation of these brain messenger RNAs by miR-139-5p is yet to be demonstrated, it is difficult to speculate on the role of these genes in the onset of the next reproductive cycle. However, several lines of evidence in other animals are consistent with a participation in this process. In Hu sheep, GRID2 (Glutamate Ionotropic Receptor Delta Type Subunit 2) was hypothesized to contribute to the high fecundity trait in a genetic screen [28]. This genetic link to reproduction was also observed in goat [29] and pig [30]. Further investigations are needed to disentangle the role of miR-139 in regulating the onset of the next reproductive cycle in iteroparous species.

### miR-202: a key miRNA for reproduction and biomarker of egg quality

miR-202 is a microRNA predominantly expressed in male and female gonads in vertebrates. miR-202-5p was originally identified as predominantly expressed in ovary, testis and uterus in mice (*Mus musculus*) [31]. Subsequent analyses in frog (*Xenopus tropicalis*) showed that miR-202-5p was highly abundant in the female germline but could not be detected in a number of adult somatic tissues including muscle, heart, brain, and liver [32]. In fish, miR-202 has been reported as predominantly expressed in halibut (*Hippoglossus hippoglossus*) gonads, compared to brain [33]. In rainbow trout, we showed that miR-202-5p was abundant in both testis and ovary, while not detected in a number of adult somatic tissues including gills, brain, pituitary, liver, intestine, stomach, heart, muscle, spleen, trunk kidney and head kidney [8, 34]. In rainbow trout, miR-202 accounts for 20 and 10 % of the total reads in testis and ovary, respectively [13]. This remarkable gonad-predominant pattern is also observed in medaka (*Oryzias latipes*) [10], zebrafish (*Danio rerio*) [12, 35] and three-spined stickleback (*Gasterosteus aculeatus*) [12]. RNA-seq data available in the FishmiRNA database [17] also clearly highlight the strong gonad-predominant expression of miR-202-5p in two holostean (i.e. the sister group of teleosts) species, the spotted gar (*Lepisosteus oculatus*) and the bowfin (*Amia calva*), and in a number of teleosts including Allis shad (*Alosa alosa*), striped catfish (*Pangasianodon hypophthalmus*), black bullhead (*Ameiurus melas*), Zebrafish, Eastern mudminnow (*Umbra pygmae*), three-spined stickleback, medaka, and European perch (*Perca fluviatilis*). In all fish species investigated to date, including rainbow trout as well as holostean and teleostean species available in the FishmiRNA database, miR-202-5p is by far the most abundant mR-202 mature form. This indicates that miR-202-5p is the biologically active miR-202 mature form, while miR-202-3p is the minor/passenger form. The very strong and evolutionarily-conserved gonad-predominant expression of miR-202 is associated with difference in miRNA levels at different steps of the reproductive cycle in the rainbow trout ovary [34]. In rainbow trout, miR-202-5p levels in blood plasma exhibit a dramatic increase at the time of ovulation, compared to previtellogenic and vitellogenic periods [8]. Together, these observations suggest an important role for miR-202 in the reproductive physiology of fishes, and possibly vertebrates. In the medaka, functional analyses have shown that miR-202 was involved in the regulation of reproductive cycle in both males and females [10]. Subsequent functional *in vivo* study revealed that miR-202 was regulating fecundity by targeting *tead3b*, a member of the Hippo signaling pathway [11]. In the present study, we observed a strong down-regulation of miR-202-5p levels in ovarian fluid after ovulation, when eggs are retained in the body cavity. This pattern is fully consistent with a role in ovarian function during or prior to ovulation in rainbow trout.

In addition of being a key miRNA for reproduction, our results also highlight the potential of miR-202 to serve as a non-invasive biological marker of egg quality. The identification of egg quality biomarkers is a long-standing goal in aquaculture of for wild-species restauration purposes. To date, there is currently no accurate marker of egg quality in salmonids. Here we show that miR-202-5p levels in ovarian fluid clearly can be used to discriminate high quality eggs sampled at ovulation from low quality eggs sampled 21 days after ovulation. Importantly the non-invasiveness nature of ovarian fluid sampling through gentle stripping under anesthesia is especially interesting.

## Materials and methods

### Experimental design and biological fluid sampling

To analyze the evolution of blood plasma and ovarian fluid c-miRNAome in relationship with the decrease of egg quality occurring over time following ovulation, fish were sampled at ovulation and, again, 21 days post-ovulation. This was carried out in parallel in two groups of six females held in 100-liter tanks. In one group (eggs retained), eggs were left in the body cavity, while in the other group (eggs removed), eggs were removed from the body cavity upon ovulation. During the pre-ovulatory period (i.e. before the beginning of sampling), females were checked under anesthesia twice a week to detect ovulation. The first sampling day corresponded to the time of detected ovulation. Blood samples (2 ml) were taken from all females at ovulation and 21 days after ovulation. Blood samples were collected from the caudal vein using EDTA-coated syringes (sodium EDTA, 10%). The blood samples were centrifuged (3000 g, 15 min, 4°C), and the plasma samples were aliquoted, frozen in liquid nitrogen, and stored at −80°C until further analysis. Ovarian fluid samples (2 ml) were collected at ovulation from all females and, again, 21 days after ovulation from the females of the ‘eggs retained’ group. Ovarian fluid was collected after manual stripping of a small amount the eggs from ovulated females over a mesh screen. The collected ovarian fluid was then centrifuged (3000 g, 15 min, 4°C) to pellet cells and debris, aliquoted, frozen in liquid nitrogen, and stored at −80°C until further analysis.

For the histological analysis, fish were sampled at ovulation and approximately three weeks after ovulation. Ovulation was checked 2-3 times a week as described above. Eggs were either manually stripped at the time of detected ovulation (eggs removed) or not (eggs retained). For sampling, fish were euthanized and the ovary was dissected out of the body cavity.

### Histology and image analysis

For histological analysis, ovaries were dissected from euthanized females, snap-frozen in liquid nitrogen, and stored at –80 °C until further processing. The ovaries were then fixed in 4% paraformaldehyde (PFA) at 4 °C overnight, dehydrated in 100% methanol, and stored at –20 °C. Fixed ovaries were embedded in paraffin wax and sectioned at a thickness of 7 µm using a microtome (HM355, Microm). Only median sections of the ovaries were retained for further analysis. The nuclei in the median sections were stained with DAPI (0.1 µg/ml) at room temperature in the dark for 15 minutes. Sections were then washed in PBS overnight at 4 °C. Whole-section images were acquired using a NanoZoomer scanner (Hamamatsu). For quantitative image analysis, the area of individual oocytes was measured using Visilog 7.2 image analysis software (Windows). Based on DAPI staining intensity, nuclei were segmented, and the area enclosed by each nucleus, corresponding to an individual oocyte, was measured.

### RNA preparation

Blood plasma and ovarian fluid samples were homogenized in Trizol reagent at a ratio of 400 μL of fluid per milliliter of reagent and total RNA was extracted according to manufacturer’s instructions. During RNA extraction, glycogen was added to each plasma sample to facilitate visualization of precipitated RNAs [8].

### Small RNA sequencing

Illumina sequencing libraries were constructed using the NEXTflex small RNA kit v3 (Bioo Scientific). Starting from 1 μg of total RNA, an adapter was ligated on the 3ʹ end of the small RNAs. A second adapter was ligated to the 5ʹ end. Ligated small RNAs were subjected to reverse transcription using M-MuLV transcriptase and a RT primer complementary to the 3ʹ adapter. PCR amplification (16 cycles) was performed on the cDNA using a universal primer and a barcoded primer. Final size selection was performed on 3% gel cassette on a Pippin HT between 126pb and 169pb. Sequencing (single read 50 nucleotides) was performed using a HiSeq6000 (Illumina) with SBS (Sequence By Synthesis) technique. After quality filter, 309,323,388 million reads were obtained with a number of read per library ranging from 1,014,867 to 12,433,091 million (mean=7,364,842). Raw reads were deposited into NCBI Sequence Read Archive under BioProject accession # PRJNA693437. Reads were trimmed of the adaptor sequence GCCTTGGCACCCGAGAATTCCA and of the random primers using Cutadapt [36].

### sRNA-seq analysis

The rainbow trout miRNAome was annotated using Prost! [12], which was run on the rainbow trout reference genome (NCBI RefSeq assembly accession GCF_013265735.2). Prost! was run using the rainbow trout miRNA annotation previously established [8] and the sequencing samples of the circulating fluids. Normalized miRNA counts in reads per million (RPM) were extracted from the “compressed by annotation” Prost! output tab. When Prost! identified multiple potential annotations for a given sequence, counts for this sequence were distributed in proportion to the relative abundance of single annotated sequences of the same miRNA.

Differential expression was analyzed using DESeq2 [37] and raw counts from miRNA expression in the fluids. For each differential expression test, log fold changes were corrected using lfcShrink (type = “apeglm”), and p-values were corrected using the Benjamini-Hochberg method, which considers multiple testing to decrease the false-discovery rate. To identify miRNAs that were differentially expressed among the differing conditions, a model was built using all samples and considering all possible effects. All pairs of conditions were compared, including an interaction term, and miRNA was considered to be differentially expressed when expression in at least one comparison differed. Expression in the fluid samples was visualized by performing principal component analyses (PCA) of log-transformed DESeq2 counts. To capture the greater importance of abundant miRNAs, the PCAs were centered but not scaled. PCAs were calculated using the R package FactoMineR [38].

### miRNA target prediction

We performed a comparative genomics analysis to identify miR-139-5p target genes associated with iteroparity in salmonid fish. Five species were analyzed, representing two reproductive strategies, iteroparity (i.e. species that reproduce multiple times) and semelparity (i.e. species that reproduce only once). The following iteroparous species were used: Atlantic Salmon (*Salmo salar*), brown Trout (*Salmo trutta*), and rainbow Trout (*Oncorhynchus mykiss*). We also analyzed Chinook Salmon (*Oncorhynchus tshawytscha*) and Coho Salmon (*Oncorhynchus kisutch*), two semelparous species belonging to the Oncorhynchus genus similarly to rainbow trout. miR-139-5p target predictions were performed using TargetScan [39] with the rainbow trout sequence that is conserved in the five species. The 3’ UTR regions were retrieved from Ensembl.

### Gene expression across different organs in mature and immature male and females

Tissue expression data were normalized using log2(TPM + 1) transformation. Z-scores were calculated per gene per species across tissues. Hierarchical clustering (Euclidean distance, complete linkage) was performed using ComplexHeatmap to identify tissue-specific expression patterns.

### Ethics

Experiments and procedures were fully compliant with French and European animal welfare policies and followed guidelines of the INRAE LPGP Institutional Animal Care and Use Ethical Committee, which specifically approved this study.

## Supporting information

Supplemental datafile 1

Supplemental datafile 2

Supplemental datafile 3

Supplemental datafile 4

Supplemental datafile 5

## Data accessibility

All transcriptomics datasets generated as part of the study are publicly available. RNAseq datasets are available on the NCBI SRA portal under BioProject accession # PRJNA693437 (https://www.ncbi.nlm.nih.gov/bioproject/PRJNA693437). Corresponding SRA accession # for all samples are liste below. Ovarian fluid at ovulation (SRR13488685, SRR13488686, SRR13488574, SRR13488528, SRR13488506, SRR13488517). Ovarian fluid at 21 days post-ovulation (SRR13488607, SRR13488618, SRR13488629, SRR13488640, SRR13488651, SRR13488662). Blood plasma, eggs retained, at ovulation (SRR13488585, SRR13488540, SRR13488596, SRR13488551, SRR13488562, SRR13488573). Blood plasma, eggs removed, at ovulation (SRR13488531, SRR13488532, SRR13488533, SRR13488534, SRR13488529, SRR13488530). Blood plasma, eggs retained, at 21 days post-ovulation (SRR13488522, SRR13488523, SRR13488524, SRR13488525, SRR13488526, SRR13488527). Blood plasma, eggs removed, at 21 days post-ovulation (SRR13488515, SRR13488516, SRR13488518, SRR13488519, SRR13488520, SRR13488521).

All codes and scripts used for miRNA processing are available here : https://github.com/INRAE-LPGP/phenomir.

## Author’s Contributions

Conceptualization, E.C., V.T., and J.B.; formal analysis, E.C., L.M., M.R.A., V.T., C.L. and J.B.; investigation, E.C., C.L., L.M., and J.B.; writing—original draft preparation, E.C., M.R.A., and J.B.; writing—review and editing, E.C., M.R.A., and J.B.; visualization, E.C., C.L., M.R.A., and J.B.; supervision, V.T. and J.B.; project administration, E.C. and J.B.; funding acquisition, E.C. and J.B. All authors have read and agreed to the published version of the manuscript.

## Acknowledgments

The authors thank the INRAE LPGP fish facility staff for fish breeding.

